# Endoplasmic reticulum calnexins participate in the primary root growth response to phosphate deficiency

**DOI:** 10.1101/2022.11.01.514754

**Authors:** Jonatan Montpetit, Joaquín Clúa, Yi-Fang Hsieh, Evangelia Vogiatzaki, Jens Müller, Steffen Abel, Richard Strasser, Yves Poirier

## Abstract

Accumulation of incompletely folded proteins in the endoplasmic reticulum (ER) leads to ER stress, activates ER protein degradation pathways, and upregulates genes involved in protein folding (Unfolded Protein Response; UPR). ER stress has been associated with abiotic stress conditions that affect protein folding, including salt stress. However, the role of ER protein folding in plant responses to nutrient deficiencies is unclear. We analyzed several *Arabidopsis thaliana* mutants affected in ER protein quality control and established that both *CALNEXIN* (*CNX*) genes function in the primary root’s response to phosphate (Pi) deficiency. CNX and calreticulin (CRT) are homologous ER lectins that bind to N-glycosylated proteins to promote their folding. Growth of *cnx1-1* and *cnx2-2* single mutants was similar to that of the wild type under high and low Pi conditions, but the *cnx1-1 cnx2-2* double mutant showed decreased primary root growth under low Pi conditions due to reduced meristematic cell division. This phenotype was specific to Pi deficiency; the double mutant responded normally to osmotic and salt stress. The root growth phenotype was Fe dependent and was associated with Fe accumulation in the root. Two genes involved in Fe-dependent inhibition of root growth under Pi deficiency, the ferroxidase gene *LPR1* and P5-type ATPase *PDR2*, are epistatic to *CNX1/CNX2*. Overexpressing *PDR2* failed to complement the *cnx1-1 cnx2-2* root phenotype. *cnx1-1 cnx2-2* showed no evidence of UPR activation, indicating a limited effect on ER protein folding. CNX might process a set of N-glycosylated proteins specifically involved in the response to Pi deficiency.

**One sentence summary:** Calnexin, a lectin chaperone engaged in the folding of N-glycosylated proteins in the ER, participates in primary root adaptation to low phosphate conditions.

## Introduction

The endoplasmic reticulum (ER) serves as the major entry point for proteins into the secretory pathway as well as for proteins destined for the plasma membrane (PM). It is estimated that approximately one-third of cellular proteins pass through this organelle (Strasser, 2018). The ER is thus a major site for folding and quality control of proteins involved in numerous cellular processes, including cell wall synthesis, nutrient transport, and PM-based signal transduction (Brandizzi, 2021). The ER harbors two main pathways to assist in protein folding. The first pathway involves the general chaperones BiPs, which belong to the classical heat shock protein 70 (HSP70) family, the DNA J protein ERdj3 and its associated stromal-derived factor 2 (SDF2) protein, and protein disulfide isomerases (PDI), which promote the formation of disulfide bonds (Strasser, 2018). The second pathway, a distinct ER folding pathway known as the calnexin-calreticulin cycle, is dedicated to N-glycosylated proteins. Calnexin (CNX) and calreticulin (CRT) are lectins that share a common architecture consisting of two major domains: a glycan binding domain and a long flexible P-domain involved in recruiting other co-chaperones such as PDIs. While CNX is anchored to the ER via a transmembrane domain, its homologue CRT is soluble within the ER matrix and harbors a luminal KDEL ER retrieval signal (Strasser, 2018; Kozlov and Gehring, 2020). *Arabidopsis thaliana* contains two CNX genes and three CRT genes (Persson et al., 2003; Liu et al., 2017).

In the CNX-CRT cycle, proteins entering the ER are first conjugated with a Glc_3_Man_9_GlcNAc_2_ glycan on specific asparagines by the oligosaccharyltransferase (OST) complex. The N-linked glycans are then trimmed by two glucosidases (GCSI and GCSII) to generate a monoglucosylated protein, which specifically binds to CNX or CRT to promote protein folding and maturation. Removal of the terminal glucose by GCSII leads to the release of the glycoprotein from CNX/CRT. If the protein is inappropriately folded after release, the glucosyltransferase UDP-glucose:glycoprotein glucosyltransferase (UGGT) adds back a terminal glucose, enabling the re-association of the misfolded glycoprotein with CNX or CRT and thus initiating an additional round of folding (Liu and Howell, 2010; Strasser, 2018).

ER proteins that repeatedly fail to properly fold after several rounds of the CNX-CRT cycle are directed to become degraded. An important pathway for ER protein degradation involves the translocation of misfolded proteins to the cytosol for proteasomal degradation, a process termed ER-associated degradation (ERAD). Protein degradation through ERAD involves the recognition and transport of misfolded proteins across the ER membrane to the cytosol, followed by polyubiquitination and degradation via the 26S proteasome (Chen et al., 2020). The accumulation of misfolded proteins in the ER leads to ER stress and the activation of the unfolded protein response (UPR). In turn, the activation of the UPR results in the upregulation of genes involved in vesicular trafficking, ERAD, and protein folding, including *BiPs* and *PDIs* (Liu and Howell, 2016). The UPR signaling pathway has two branches. In the first branch, the ER-anchored RNA splicing factor IRE1 modifies the mRNA of the transcription factor bZIP60, yielding a form of bZIP60 that lacks a transmembrane domain and is targeted to the nucleus. The second branch of the UPR signaling pathway activates two other members of the bZIP family, bZIP17 and bZIP28, via protease processing in the Golgi (Liu and Howell, 2016). Chronic ER stress that cannot be resolved by the activation of ERAD and the UPR can lead to programmed cell death as well as autophagy (Manghwar and Li, 2022).

ER stress has been associated with numerous abiotic stress factors that are thought to lead to defects in protein folding in the ER, such as heat, drought, osmotic, salt, and metal stress. The link between the control of ER protein folding and abiotic stress has been demonstrated via the analysis of mutants as well as transgenic plants overexpressing genes encoding ER chaperones, such as *BiP, CNX*, and *PDIs*, as well as genes involved in the ERAD and UPR pathways, including *IRE1* and *bZIP28* (Gao et al., 2008; Deng et al., 2011; Kim et al., 2013; Joshi et al., 2019; Park and Park, 2019; Reyes-Impellizzeri and Moreno, 2021). However, whether the control of protein folding in the ER has a role in plant responses to nutrient deficiency has not been determined, although recent work has shown that autophagy may be implicated in such stress (Naumann et al., 2019; Stephani et al., 2020; Yoshitake et al., 2021).

Phosphorus is one of the most important nutrients affecting plant growth in both agricultural and natural ecosystems (Poirier et al., 2022). Plants acquire phosphorus almost exclusively via the transport of soluble inorganic phosphate (H_2_PO_4_^−^; Pi) into roots. Plants have evolved a series of metabolic and developmental responses to Pi deficiency that are aimed at maximizing Pi acquisition from the environment and optimizing its internal use for growth and reproduction (Dissanayaka et al., 2021; Poirier et al., 2022). One of the best-characterized responses of roots to phosphate deficiency is a decrease in primary root growth associated with reduced root meristem size (Crombez et al., 2019). This phenotype has been associated with the presence of Fe^+3^-malate complexes in the root meristem, which generate reactive oxygen species (ROS), and in turn lead to changes in the cell wall structure and inhibition of cell-to-cell communication (Müller et al., 2015; Balzergue et al., 2017; Mora-Macias et al., 2017). Genetic screens for genes that contribute to changes in primary root growth under Pi deficiency identified *LPR1* and *LPR2*, encoding ferroxidases that convert Fe^+2^ to Fe^+3^, and *PDR2*, encoding an ER-localized P5-type ATPase thought to negatively affect LPR activity via an unknown mechanism (Ticconi and Abel, 2004; Svistoonoff et al., 2007; Ticconi et al., 2009; Naumann et al., 2022). Additional proteins found to participate in this pathway include the malate and citrate efflux channel ALMT1; the STOP1 transcription factor, which regulates *ALMT1* expression; ALS3 and STAR1, which together form a tonoplast ABC transporter complex involved in plant tolerance to aluminum (although the nature of the molecule that is transported remains to be defined); and the CLE14 peptide receptors CLV2 and PEPR2 (Balzergue et al., 2017; Dong et al., 2017; Gutierrez-Alanis et al., 2017; Mora-Macias et al., 2017).

In the present study, we analyzed Arabidopsis mutants affected in components of ER protein folding and quality control for their response to phosphate deficiency. We determined that CNX proteins participate in the Fe-dependent inhibition of primary root growth in response to phosphate deficiency.

## Results

### The *cnx1 cnx2* double mutant shows reduced primary root growth under low Pi conditions

We crossed the Arabidopsis *cnx1-1* mutant (SALK_083600), which has a T-DNA insertion in the 3^rd^ exon of *CNX1* (At5g61790), with *cnx2-2* (SAIL_865_F08) and *cnx2-3* (SAIL_580_H02), which have T-DNA insertions in the third exon of *CNX2* (At5g07340), to create two independent double mutant combinations (Figure 1A). Immunoblot analysis of protein extracts from whole seedlings showed that CNX proteins were absent in the *cnx1-1 cnx2-2* double mutant, indicating that these mutant alleles are likely null (Figure 1B). We grew the plants in fertilized soil and in clay irrigated with nutrient solution containing 1 mM Pi (high Pi; HPi) or 75 µM Pi (low Pi; LPi) and found no significant differences between the single and double mutants compared to the wild type Col-0 in terms of fresh weight (Figure 1C and D) or Pi content (Figure 1E) in roots or rosettes. By contrast, in seedlings grown on solid medium, primary root length was significantly reduced in the *cnx1-1 cnx2-2* and *cnx1-1 cnx2-3* double mutants compared to Col-0 under LPi but not HPi conditions (Figure 2A). This phenotype was complemented by transforming the *cnx1-1 cnx2-2* double mutant with the *CNX1-GFP* or *CNX2-GFP* fusion construct driven by their respective endogenous promoters (Figure 2B). Confocal microscopy of roots of the complemented lines expressing CNX1-GFP or CNX2-GFP revealed localization of these fusion proteins in the ER (Supplemental Figure S1A). Co-localization of CNX1-GFP and CNX2-GFP with an ER marker (ER-RFP) was observed in transiently transfected tobacco (*Nicotiana benthamiana*) leaf cells (Supplemental Figure S1B).

**Figure 1.**
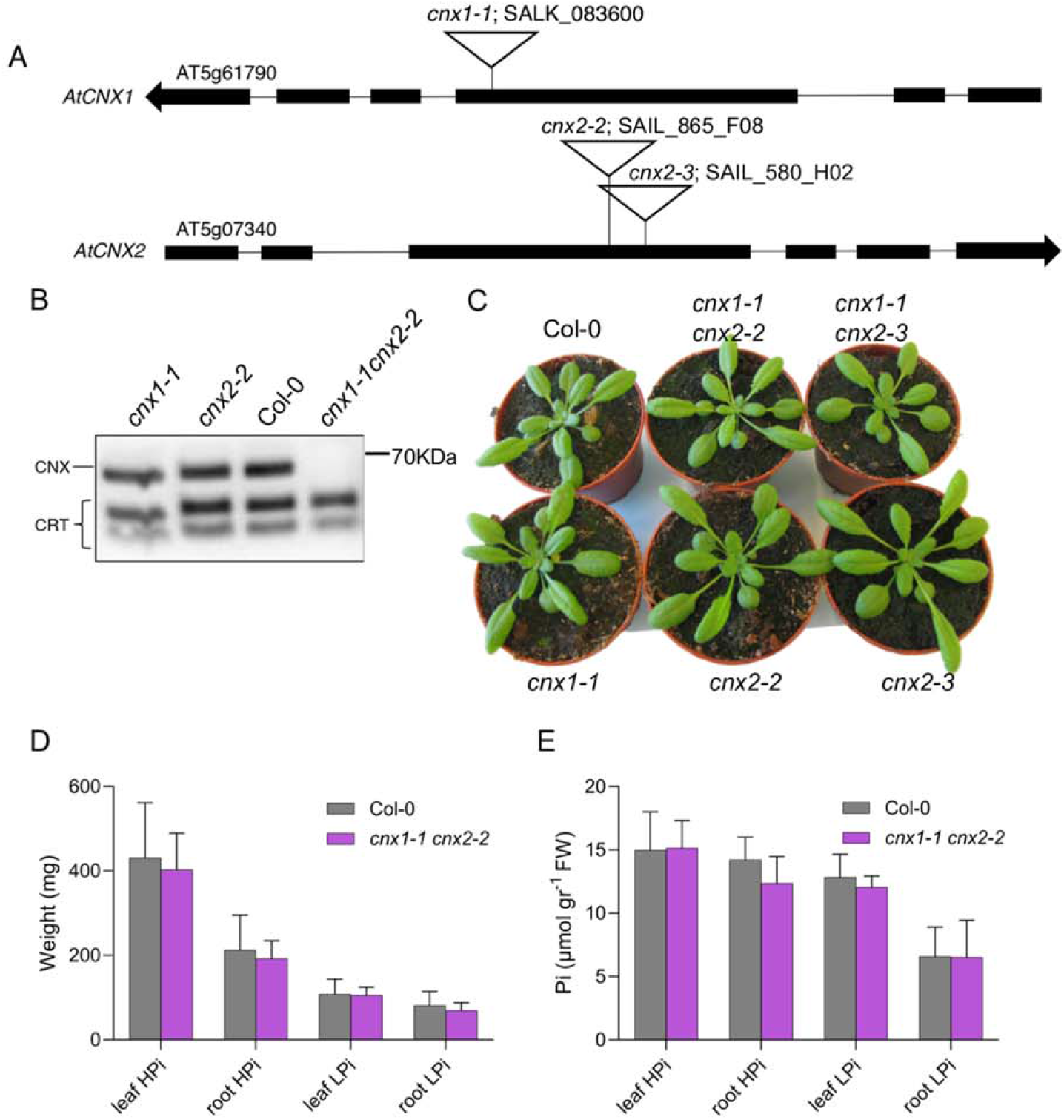
Phenotype of the *cnx1 cnx2* double mutant in soil. **(A)** Schematic diagram of the T-DNA insertions in the *CNX1* (At5g61790) and *CNX2* (At5g07340) genes in the *cnx* mutants. Exons are shown as black boxes. **(B)** Immunoblot analysis of CNX and CRT in whole protein extracts from seedlings. The position of the 70 KDa molecular weight marker is shown on the right. **(C)** Rosettes of 3.5-week-old plants grown in soil. **(D, E)** Fresh weight (D) and Pi content (E) in whole rosettes (leaf) and roots of plants grown for 4 weeks in clay irrigated with nutrient solution containing 1 mM Pi (HPi) or 75 µM Pi (LPi). Statistical analysis was performed by Student’s *t*-test compared to the Col-0 control, error bars = SD, n = 8-10.

**Figure 2.**
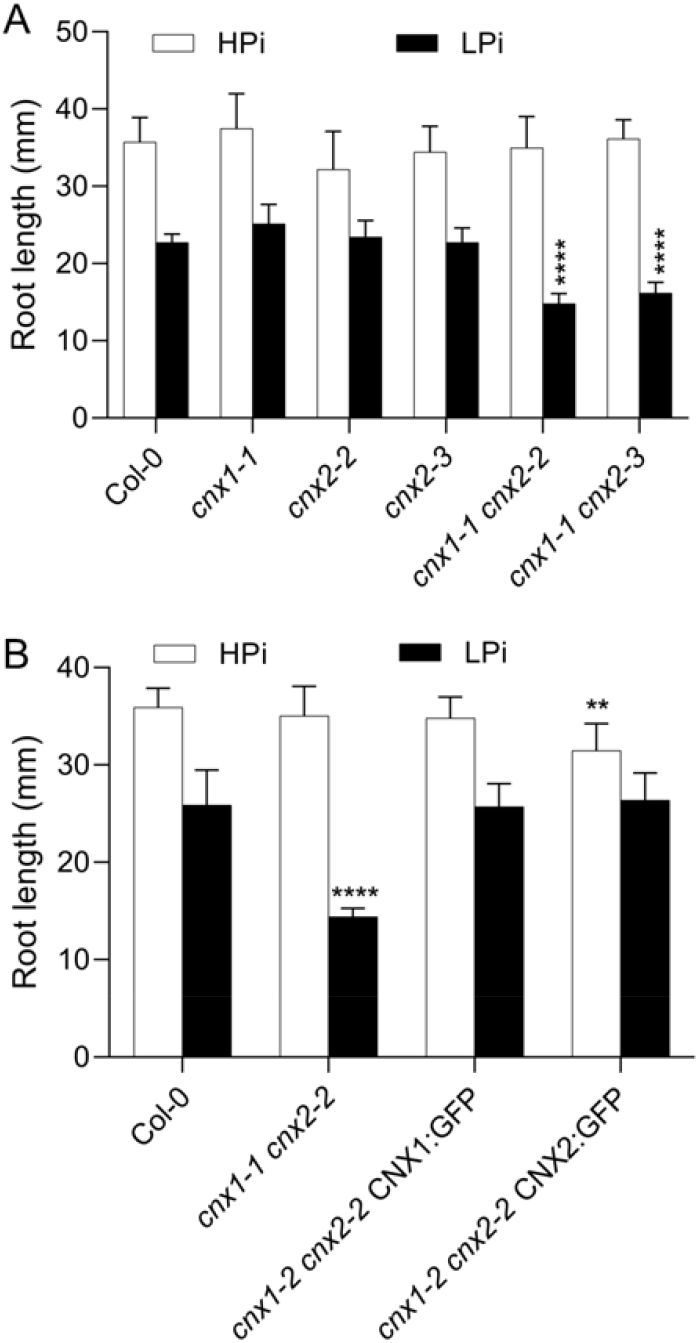
Primary root growth of the *cnx1 cnx2* double mutant under high and low Pi conditions. **(A)** Primary root length of Col-0 compared to *cnx1-1* and *cnx2-2* single and double mutants. **(B)** Complementation of the primary root phenotype of *cnx1-1 cnx2-2* plants transformed with the CNX1:GFP or CNX2:GFP construct. Plants were grown for 7 days on plates containing 1 mM Pi (HPi) or 75 µM Pi (LPi) before measuring primary root length. Statistical analysis was performed by two-way ANOVA followed by a Tukey’s test, and significant differences compared to Col-0 in each growth condition are shown: **, P < 0.01; ***, P < 0.001; ****, P < 0.0001; error bars = SD; n ≥ 9.

### Mutants in other components of the CNX/CRT cycle and ER chaperone system do not reproduce the *cnx1 cnx2* root growth phenotype under low Pi

In addition to CNX, ER protein quality control relies on numerous other proteins, including chaperones and enzymes involved in glycosylation and glycan modifications in the ER (Strasser, 2018). We therefore examined primary root growth of mutants in various components of the CNX/CRT cycle and ER protein quality control under LPi conditions. Arabidopsis CRTs are encoded by three genes, which are divided into two groups based on sequence homology and function: *CRT1/CRT2* and *CRT3* (Persson et al., 2003; Christensen et al., 2010). No significant differences were detected in the root growth of *crt1 crt2* or *crt3* mutants under HPi or LPi conditions compared to Col-0 (Figure 3A).

**Figure 3.**
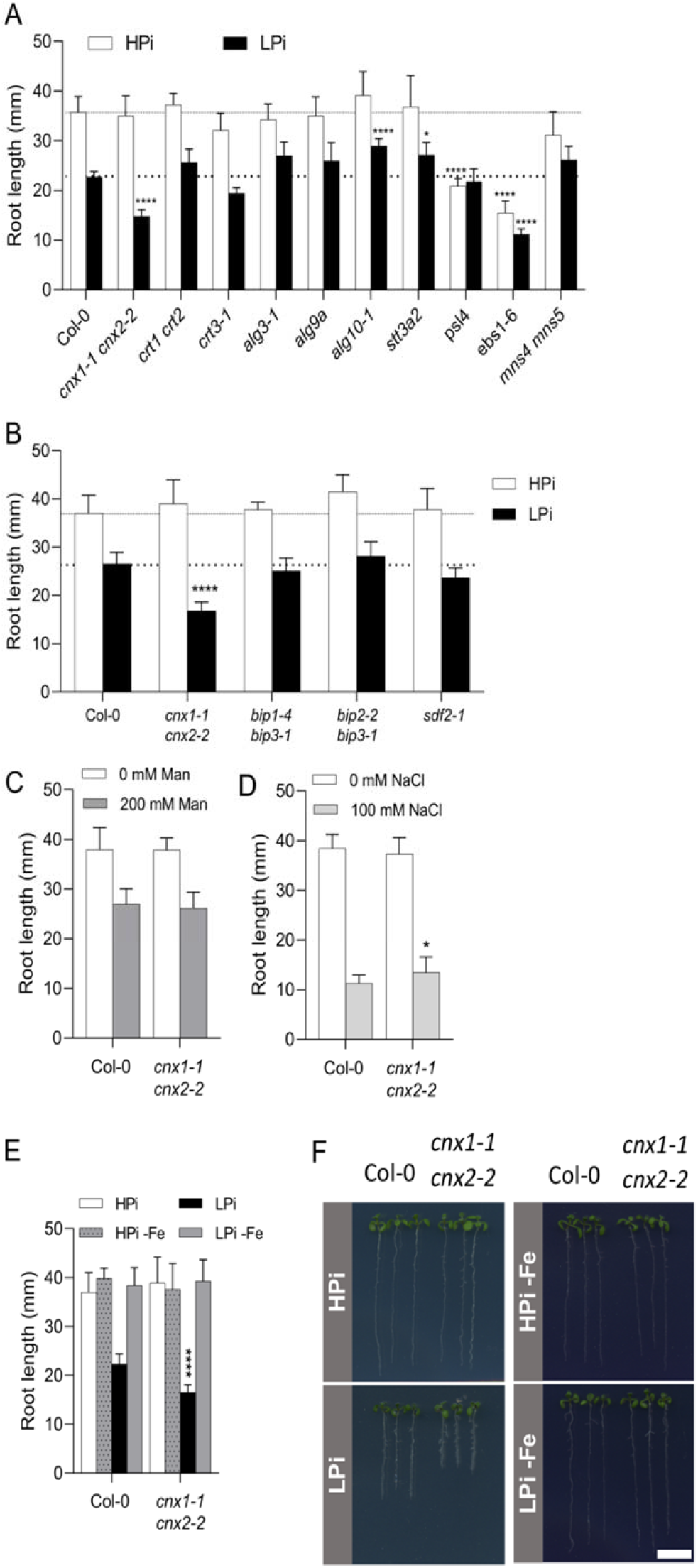
Primary root growth of mutants in genes involved in ER protein synthesis and quality control. **(A-B)** Plants were grown for 7 days on plates containing HPi or LPi before measuring primary root length. **(C-D)** Primary root length of Col-0 and *cnx1-1 cnx2-2* plants after 7 days of growth on HPi plates (C) without or with 200 mM mannose or (D) without or with 100 mM NaCl. **(E-F)** Primary root length of Col-0 and *cnx1-1 cnx2-2* after 7 days of growth on plates containing HPi or LPi half-strength MS medium or the same medium with ferrozine to chelate Fe (HPi-Fe and LPi-Fe). Statistical analysis was performed by two-way ANOVA followed by a Tukey’s test, and significant differences compared to Col-0 in each growth condition are shown, *P < 0.05, **P < 0.01, ***P < 0.001, ****P < 0.0001, error bars = SD, n ≥ 5. Bar represents 1 cm in F.

The synthesis of the core oligosaccharide unit Glc_3_Man_9_GlcNAc_2_ involves a series of ER glycosyltransferases including the mannosyltransferases ALG3 and ALG9 and the glucosyltransferase ALG10 (Kajiura et al., 2010; Farid et al., 2011; Hong et al., 2012). Following its synthesis, the Glc_3_Man_9_GlcNAc_2_ unit is added to ER proteins co-translationally by the membrane-associated heteromeric OST complex, which includes the catalytic STT3 subunit encoded by *STT3A* in Arabidopsis (Koiwa et al., 2003). Primary root growth under HPi and LPi conditions was not reduced in *alg3-1, alg9a, alg10-1*, or *stt3a2* mutants compared to Col-0 (Figure 3A).

The presence of terminal α1,2-linked glucose residues, which facilitate the interaction between CNX/CRT and N-glycosylated proteins, is regulated by the trimming action of GCSII and the glucosylating action of UGGT. *PSL4* encodes the β-subunit of GCSII (Lu et al., 2009). The primary roots of the *psl4* mutant were shorter than Col-0 when grown on Hpi medium, but there was no significant further reduction in their length when grown on LPi medium (Figure 3A). Primary root growth was severely compromised in the *ebs1-6/uggt1-1* mutant on HPi medium, and this effect was only slightly enhanced on LPi medium (Figure 3A).

ER proteins that pass through the CNX/CRT cycle but remain inappropriately folded are degraded by ERAD. This process involves the trimming of mannosyl groups on the N-glycan chain by the α-mannosidases MNS4 and MNS5 (Huttner et al., 2014). Primary root growth of the *mns4 mns5* double mutant was not significantly different from Col-0 on HPi or LPi medium (Figure 3A).

We also examined the role of the ER chaperone pathway involving BiP and SDF2 in the response of Arabidopsis roots to Pi deficiency. While SDF2 is encoded by a single gene in Arabidopsis (Nekrasov et al., 2009), three genes encode the ER BiP chaperones. *BIP1* and *BIP2* encode proteins that are 99% identical and are ubiquitously expressed, while the more divergent *BiP3* is expressed under ER stress (Maruyama et al., 2014). Root growth of the *bip1-4 bip3-1, bip2-2 bip3-1*, and *sdf2-1* mutants was similar to that of Col-0 on both HPi and LPi media (Figure 3B).

Several mutants related to the CNX/CRT cycle and ER protein homeostasis, including *alg10, stt3a, mns4 mns5*, and *ebs1-6*/*uggt1*, exhibit strong root growth phenotypes under salt stress (Koiwa et al., 2003; Farid et al., 2011; Huttner et al., 2014; Blanco-Herrera et al., 2015). To investigate whether the reduced primary root length observed in *cnx1-1 cnx2-2* was specific to Pi deficiency stress, we examined root growth in this double mutant under two other abiotic stress conditions that reduced primary root growth: osmotic stress (200 mM mannitol) and salt stress (100 mM NaCl). Under both stress conditions, primary root growth was similar in the *cnx1-1 cnx2-2* double mutant and Col-0 (Figure 3C-D), indicating that the root growth phenotype of this double mutant is specific to Pi deficiency stress.

### The root phenotype of *cnx1 cnx2* is due to reduced root apical meristem activity

Reduced primary root growth under stress conditions can be caused by reduced cell division within the meristem, reduced cell elongation, or both. Under LPi but not HPi conditions, the meristematic zone was smaller in *cnx1-1 cnx2-2* compared to Col-0 and the corresponding single mutants (Figure 4A, B). By contrast, the cell length in the elongation zone was not significantly different between the mutants and Col-0 under HPi or LPi conditions (Figure 4A, C). These data indicate that *cnx1-1 cnx2-2* is mainly affected in meristematic cell division under LPi conditions. To further evaluate the contribution of cell division to the mutant phenotype, we introduced into the *cnx1-1 cnx2-2* double mutant a reporter construct for cell division consisting of labile GUS under the control of the cyclin B1 promoter (Colon-Carmona et al., 1999). The number of dividing, GUS-expressing cells was similar in *cnx1-1 cnx2-2* vs. Col-0 roots under HPi conditions (Figure 4D). By contrast, a clear reduction in GUS-expressing cells was observed in Col-0 roots grown under LPi, in accordance with the known reduction in meristematic cell division under these conditions (Ticconi et al., 2004). Importantly, a further reduction in GUS expression in roots was observed in the *cnx1-1 cnx2-2* double mutant compared to Col-0 on LPi (Figure 4D). Altogether, these data indicate that the altered primary root growth of *cnx1-1 cnx2-2* is primarily due to reduced meristematic cell division under LPi conditions.

**Figure 4.**
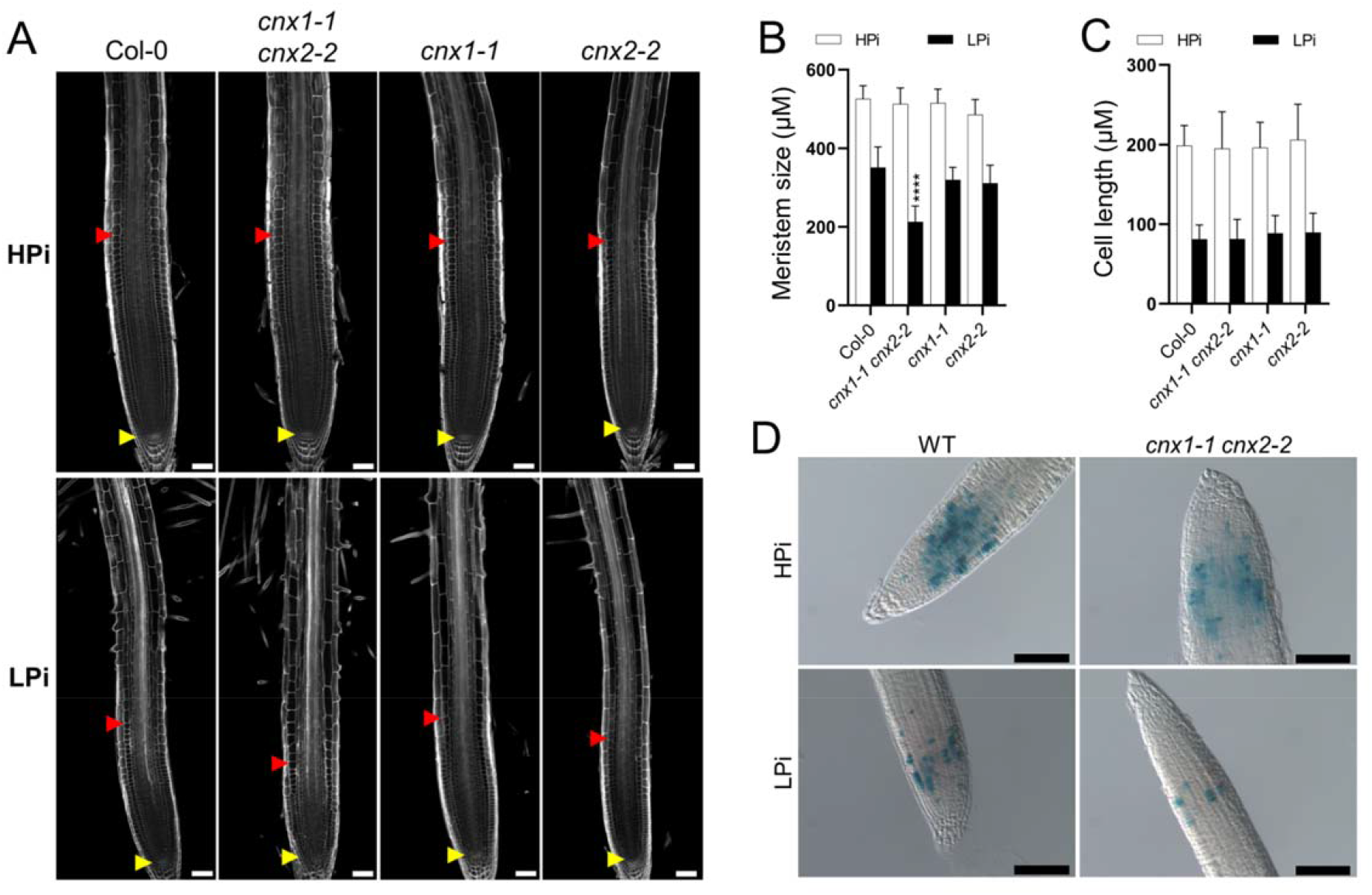
The *cnx1-1 cnx2-2* double mutant is affected in meristem activity. **(A-C)** Plants were grown for 7 days on plates containing HPi or LPi before measuring the length of the cell division zone in the meristem, defined in A by the red and red arrows (A, B) and cell length in the differentiation zone (C). Statistical analysis (B, C) was performed by two-way ANOVA followed by a Tukey’s test; significant differences compared to Col-0 under each growth condition are shown: ****, P < 0.0001; error bars = SD; n ≥ 5 in (B) and 20 in (C). **(D)** Col-0 and *cnx1-1 cnx2-2* plants transformed with the cylinB1:GUS reporter gene construct were grown for 7 days on plates containing HPi or LPi medium and stained for β-glucuronidase activity. Bars represent 50 um in A and 100 µm in D.

### The root phenotype of *cnx1-1 cnx2-2* is dependent on Fe and associated with increased Fe deposition in the meristem

Several studies have shown that the reduced primary root growth of plants under low Pi in Col-0 and in various mutants with more severe root growth inhibition is dependent on the presence of Fe in the growth medium (Ticconi et al., 2009; Müller et al., 2015; Balzergue et al., 2017; Dong et al., 2017). Indeed, a comparison of root growth on HPi and LPi medium with and without Fe showed that the reduced primary root growth observed in *cnx1-1 cnx2-2* under LPi conditions was also dependent on the presence of Fe in the medium (Figure 3E-F). We used Perls-DAB staining to examine the distribution of apoplastic Fe in plants grown under HPi and LPi conditions. The *lpr1* mutant (which is insensitive to low Pi-induced root growth inhibition) and *pdr2* (which shows very strongly reduced primary root growth under low Pi conditions) were used as controls (Müller et al., 2015). In plants grown under HPi conditions, no significant differences were observed in Fe distribution in the root meristematic and elongation zones between Col-0 and *cnx1-1 cnx2-2* or *pdr2*, whereas *lpr1* showed substantially reduced Fe deposition (Figure 5, upper panels). Under LPi conditions, the *cnx1-1 cnx2-2* double mutant showed robust enhancement of Fe deposition in the root differentiation zone and more modest enhancement in the root differentiation and meristematic zones compared to Col-0, whereas *pdr2* roots showed extensive Fe deposition throughout the root, and *lpr1* showed minimal Fe deposition (Figure 5, lower panels).

**Figure 5.**
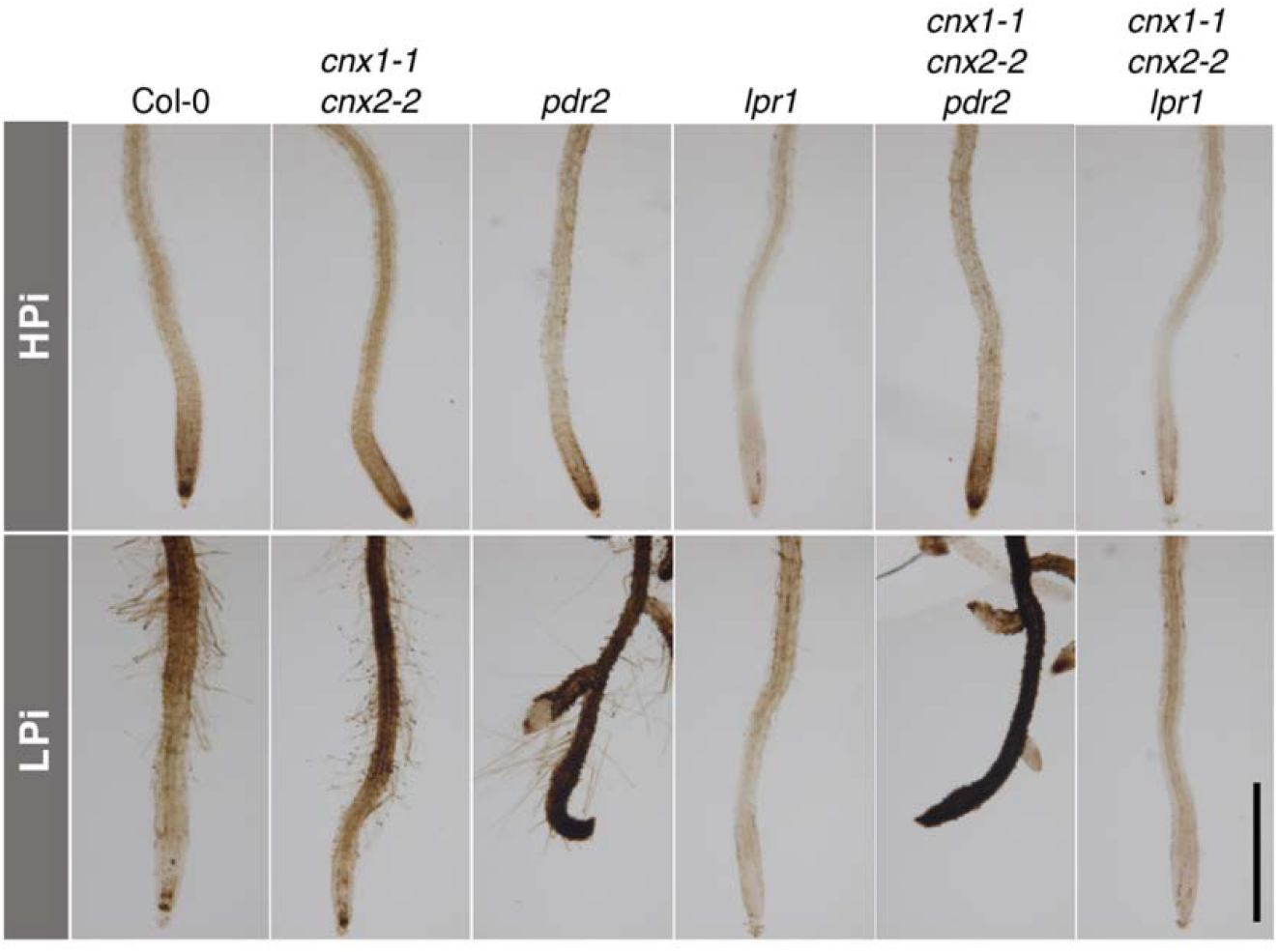
Fe accumulation and distribution in the roots of mutants grown under high and low Pi conditions. Plants were grown for 7 days on plates containing 1 mM or 75 µM Pi and subjected to Perls-DAB staining for Fe visualization. Bar represents 1 mm.

### *pdr2* and *lpr1 lpr2* are epistatic to *cnx1-1 cnx2-2*

We examined the epistasis among *cnx1-1 cnx2-2, lpr1 lpr2*, and *pdr2* by generating triple and quadruple mutants. Primary root growth of *cnx1-1 cnx2-2 lpr1 lpr2* was insensitive to low Pi, as the primary root length of this quadruple mutant was identical to that of *lpr1 lpr2* and longer than that of Col-0 under LPi conditions (Figure 6A). The *pdr2* mutant showed reduced primary root growth in HPi; this phenotype remained unchanged in the *cnx1-1 cnx2-2 pdr2* triple mutant. On LPi medium, the *pdr2* mutant showed more strongly reduced primary root growth than *cnx1-1 cnx2-2*, and this phenotype was maintained in the *cnx1-1 cnx2-2 pdr2* triple mutant (Figure 6B). The epistatic action of *lpr1* and *pdr2* over *cnx1-1 cnx2-2* was also observed at the level of Fe accumulation for roots grown under HPi and LPi (Figure 5).

**Figure 6.**
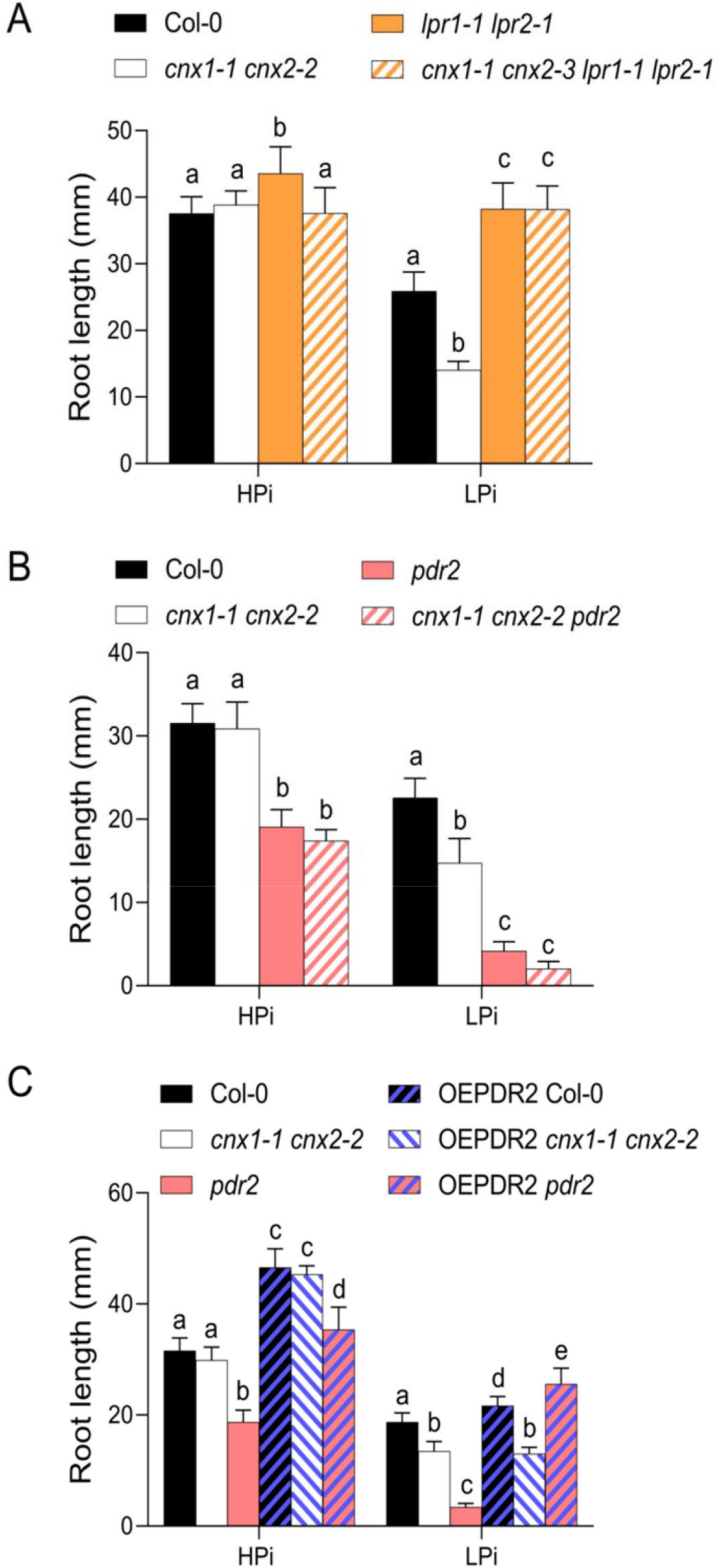
Epistatic interactions among *cnx1-1 cnx2-2, lpr1-1 lpr2-1*, and *pdr2*. Plants were grown for 7 days on HPi or LPi plates before recording primary root length. **(A)** Epistatic interaction between *cnx1-1 cnx2-2* and *lpr1-1 lpr2-1*. **(B)** Epistatic interaction between *cnx1-1 cnx2-2* and *pdr2*. **(C)** A T-DNA cassette for *PDR2* overexpression under the control of the CaMV35S promoter (OEPDR2) was introgressed into Col-0, *cnx1-1 cnx2-2*, and *pdr2*. Statistical analysis was performed by two-way ANOVA followed by a Tukey’s test, and significant differences within each growth condition are shown. Different lowercase letters (a, b, c, or d) indicate a significant difference with a P-value < 0.05, n ≥ 6.

We also examined the effect of overexpressing *PDR2* driven by the CaMV35S promoter. In both Col-0 and *pdr2*, overexpression of *PDR2* led to significantly longer primary roots compared to Col-0 plants on both HPi and LPi media. By contrast, while overexpressing *PDR2* in the *cnx1-1 cnx2-2* double mutant background also resulted in longer primary roots compared to Col-0 grown under HPi conditions, the same plants showed shorter primary roots than Col-0 and comparable root length to the *cnx1-1 cnx2-2* double mutant when grown under LPi conditions (Figure 6C). Overall, these data indicate that the primary root phenotypes of *pdr2* and *lpr1 lpr2* are epistatic to *cnx1-1 cnx2-2* under LPi and that overexpressing *PDR2* failed to rescue the short root phenotype of *cnx1-1 cnx2-2* under LPi.

### Pi deficiency induces *CNX* gene expression and ER stress

We examined the expression of *CNX1* and *CNX2* in the shoots and roots of plants grown on LPi and HPi media via quantitative RT-PCR. The expression of both *CNX1* and *CNX2* significantly increased under Pi-deficient conditions (Figure 7A). However, the increase in expression for these genes was moderate compared to that of other Pi deficiency–responsive genes, such as *MGD3* and *PHT1;4* (Figure 7B).

**Figure 7.**
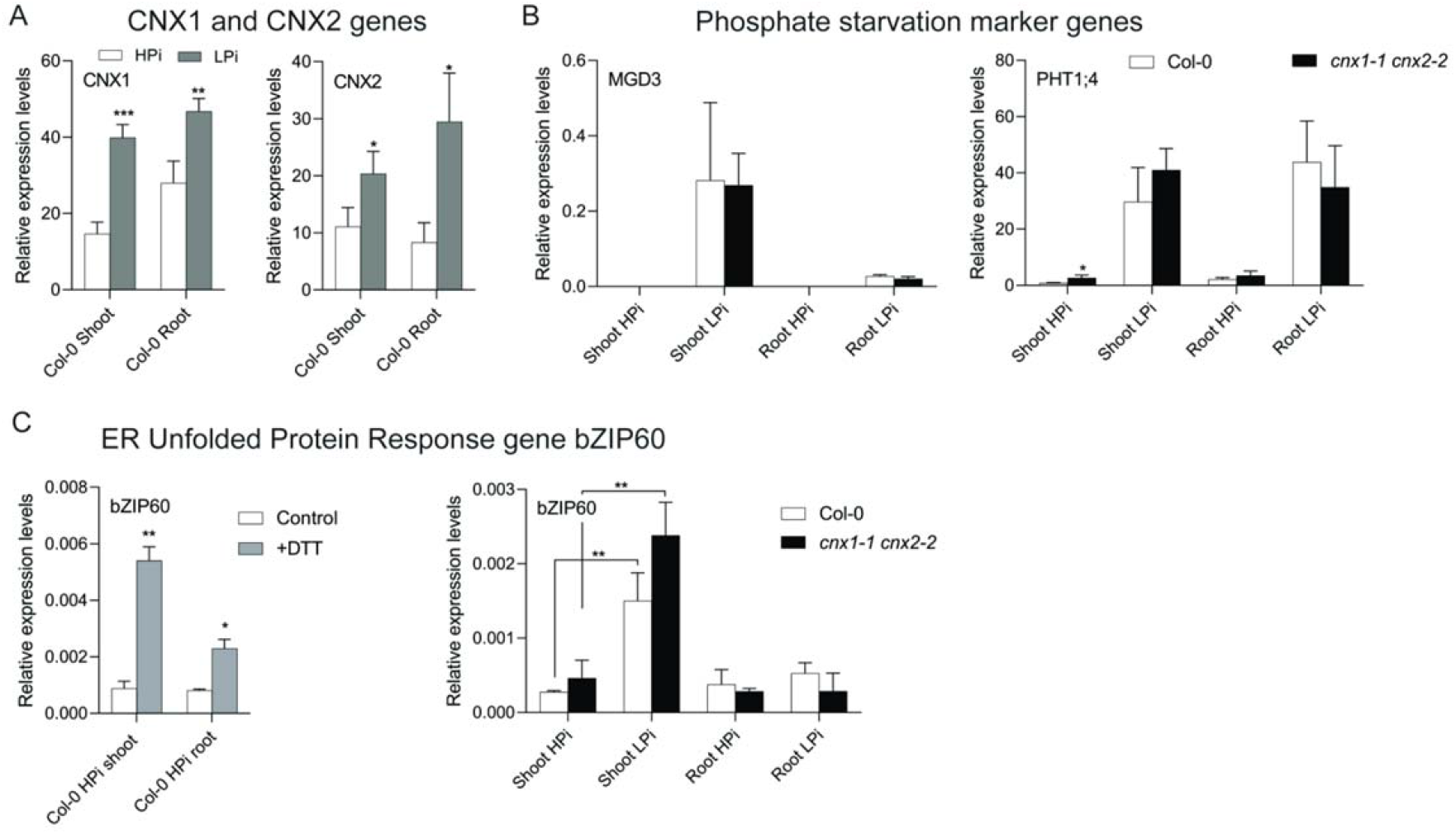
Impact of the *cnx1-1 cnx2-2* mutations on the expression of Pi deficiency and unfolded protein response marker genes. **(A)** *CNX1* and *CNX*2 expression in the shoots and roots of plants grown for 7 days in HPi or LPi medium. (B) Expression of the Pi deficiency markers *MGD3* and *PHT1;4* in the shoots and roots of Col-0 and *cnx1-1 cnx2-2* grown for 7 days on HPi or LPi medium. (C) Induction of ER Unfolded Protein Response marker gene *bZIP60* in the shoots and roots of Col-0 at 24 h after the addition of 2 mM DTT and in the *cnx1-1 cnx2-2* double mutant compared to Col-0 grown under HPi or LPi conditions. Statistical analysis was performed by Student’s *t*-test comparing different treatments (HPi and LPi for A and C, Control and DTT for C) and Col-0 vs. *cnx1-1 cnx2-2* (B, C), with significant differences indicated by asterisks:*, P < 0.05; **, P < 0.01; ***, P < 0.001. Error bars = SD, n = 3.

We investigated the transcriptional response of the *cnx1-1 cnx2-2* double mutant to Pi deficiency conditions by examining *MGD3* and *PHT1;4* expression. The expression of both genes in shoots and roots did not significantly differ between *cnx1-1 cnx2-2* and Col-0 on HPi or LPi medium, except that *PHT1;4* was slightly upregulated in *cnx1-1 cnx2-2* shoots on HPi medium (Figure 7B).

The accumulation of mis-folded proteins in the ER leads to ER stress and the increased expression of the transcription factor gene *bZIP60* (Lu and Christopher, 2008). To determine whether LPi treatment leads to ER stress and whether the *cnx1-1 cnx2-2* double mutant exhibits greater signs of ER stress compared to Col-0 plants, we compared the expression of *bZIP60* in *cnx1-1 cnx2-2* vs. Col-0 plants grown on HPi and LPi. When we treated plants with the reducing agent dithiothreitol (DTT) to induce ER stress, *bZIP60* was upregulated, with a greater increase in shoots compared to roots (Figure 7C) (Lu and Christopher, 2008). Under LPi conditions, *bZIP60* expression significantly increased in shoots but not roots in both Col-0 and *cnx1-1 cnx2-2*, with no significant difference in *bZIP60* expression between these lines (Figure 7C). Thus, the removal of calnexin did not lead to an increase in ER stress compared to Col-0 under either HPi or LPi conditions.

## Discussion

Both CNX1 and CNX2 are localized to the ER in Arabidopsis, and the corresponding genes are broadly expressed in most tissues (except that only *CNX1* is significantly expressed in pollen) and throughout development in both shoots and roots (Liu et al., 2017). Previous analysis of higher-order Arabidopsis mutants of *CNX* and *CRT* revealed that while the *cnx1 cnx2* double mutant had no phenotype under normal growth conditions, the *crt1 crt2* double mutant and the *crt1 crt2 crt3* triple mutant showed reduced rosette growth in soil and reduced hypocotyl elongation in the dark (Christensen et al., 2010; Kim et al., 2013; Vu et al., 2017). The *cnx1 crt1 crt2 crt3* quadruple mutant showed stronger defects in shoot and root developmental under normal conditions, as well as compromised fertility due to strongly reduced pollen viability and pollen tube growth. The inactivation of all *CNX* and *CRT* genes in the quintuple *cnx1 cnx2 crt1 crt2 crt3* mutant was lethal (Vu et al., 2017). Interestingly, the phenotype of the *cnx1 crt1 crt2 crt3* mutant was fully complemented by expressing either *CNX1* or *CNX2* under the control of the *CNX1* promoter, highlighting the functional redundancy of *CNX1* and *CNX2* (Vu et al., 2017).

In the present study, while the *cnx1-1* and *cnx2-2* single mutants showed no defect in primary root growth under HPi or LPi conditions, the *cnx1-1 cnx2-2* double mutant showed reduced primary root growth under LPi but not HPi conditions; this phenotype was complemented by the expression of either *CNX1* or *CNX2* driven by their native promoters. Thus, *CNX1* and *CNX2* are both required and play functionally redundant roles in the response of primary roots to Pi deficiency. Interestingly, no other mutant analyzed that is impaired in various aspects of N-glycan synthesis and the CNX-CRT cycle showed defects in primary root growth specifically under LPi conditions. These results could potentially reflect the presence of genetic redundancy or the induction of compensatory mechanisms in these mutants that do not function in the *cnx1-2 cnx2-2* mutant.

Several mutants tested for primary root growth under LPi conditions, including *alg10-1, stt3a2, ebs1-6*, and *mns4 mns5*, were previously shown to have altered root growth under salt stress (Koiwa et al., 2003; Farid et al., 2011; Huttner et al., 2014; Blanco-Herrera et al., 2015). Considering that the growth of *cnx1-1 cnx2-*2 roots was comparable to Col-0 under salt stress and osmotic stress, it is likely that defects in different components of the CNX-CRT cycle affect distinct N-glycosylated proteins to different extents. That is, the proteins affected in the *alg10-1, stt3a2, ebs1-6*, and *mns4 mns5* mutants are involved in the salt stress response, and those affected in *cnx1-1 cnx2-2* are involved in the Pi deficiency response.

The *cnx1 cnx2* mutant shares several features with the *pdr2, als3*, and *star1* mutants in terms of their responses to LPi conditions, including Fe-dependent reduced primary root growth associated with a reduction in root meristem size (Ticconi et al., 2004; Müller et al., 2015; Dong et al., 2017). However, the *pdr2, als3*, and *star1* mutants have additional root phenotypes under LPi conditions that are not observed in *cnx1-1 cnx2-2*, such as reduced cell length in the root elongation zone and a generally more distorted cellular organization in the root meristem. Furthermore, the apparent apoplastic Fe accumulation (as visualized by Perls-DAB staining) in *pdr2, als3*, and *star1* roots grown in LPi is higher in both the elongation and meristematic zones compared to *cnx1-1 cnx2-2* (Ticconi et al., 2004; Müller et al., 2015; Dong et al., 2017). Initial characterization of mutants such as *pdr2, lpr1, almt1*, and *als3* linked strong apoplastic Fe staining in the root meristematic and elongation zones with inhibited cell division and cell elongation. Fe accumulation in the meristem is associated with ROS production, which affects cell wall structure and meristem cell division via reduced mobility of SHORT-ROOT (SHR) in the stem cell niche (Müller et al., 2015; Balzergue et al., 2017). However, a more detailed analysis of dynamic changes in Fe accumulation and primary root growth over time revealed that the extent of primary root growth inhibition cannot simply be directly linked to the level of apoplastic Fe accumulation in the root meristem and elongation zone (Wang et al., 2019).

*PDR2* encodes a member of the eukaryotic type V subfamily (P5) of P-type ATPase (Ticconi et al., 2009). PDR2 is abundant in the ER, but its mode of action and transport activity are largely unknown, although recent work has reported a role of the yeast P5A ATPase Spf1 in protein translocation in the ER (McKenna et al., 2020). PDR2 is thought to modulate the activity and/or abundance of the ferroxidase LPR1 in the apoplast, which is responsible for the oxidation of Fe^+2^ to Fe^+3^ (Müller et al., 2015; Naumann et al., 2022). Consequently, the *lpr1* phenotypes (in terms of both Fe deposition and reduced primary root growth under LPi conditions) are epistatic to *pdr2* (Ticconi et al., 2009). The *lpr1* phenotypes are also epistatic to *cnx1-1 cnx2-2*. It is unknown if PDR2 is N-glycosylated and if it enters the CNX-CTR cycle. However, considering the milder phenotypes of *cnx1-1 cnx2-2* compared to *pdr2* and the finding that overexpressing *PDR2* did not influence the reduced primary root growth of *cnx1 cnx2* on LPi medium, it is unlikely that the root growth phenotype of *cnx1-1 cnx2-2* is mediated by reduced PDR2 activity.

The lack of calnexin leads to a range of phenotypes in fungi and animals, from lethality in the yeast *Schizosaccharomyces pombe* to developmental and neurological abnormalities in zebrafish, mouse, and Drosophila (Parlati et al., 1995; Kraus et al., 2010; Hung et al., 2013; Xiao et al., 2017). The current study highlights a novel role for calnexin in the response of primary root growth to Pi deficiency. Phosphate deficiency has been associated with an increase in autophagy in root tips and leaves as well as an increase in *CNX1* and *BiP2* expression (Naumann et al., 2019; Yoshitake et al., 2021). Here, Pi deficiency resulted in the increased expression of *CNX1* and *CNX2* in both roots and shoots as well as *bZIP60* in shoots. Collectively, these data reveal that Pi deficiency is associated with an increase in ER stress. Yet, the absence of a significant difference in *bZIP60* expression between Col-0 and the *cnx1-1 cnx2-2* double mutant indicates that the absence of calnexin in Arabidopsis does not lead to a systematic increase in ER stress, at least under HPi or LPi conditions. This implies that the folding and activity of a restricted number of N-glycosylated proteins are likely affected by the absence of calnexin; one or a few of these proteins likely contribute to the reduced primary root growth under LPi conditions.

A study of leucine-rich repeat receptor kinases involved in innate immunity revealed markedly different impacts of N-glycosylation on homologous receptor activity. For example, while the activity of ERF (involved in binding the bacterial elongation factor EF-Tu) was compromised by mutation of only a few of its N-glycosylation sites, the activity of the homologous protein FLS2 (a flagellin receptor) was not disrupted by mutation of several of its N-glycosylation sites (Sun et al., 2012). In accordance with these results, the activity of ERF but not of FLS2 was compromised in several mutants of genes involved in the CNX-CRT cycle, including *CRT3, SDF2, PSL4, PSL5, EBS1*, and *OST3/6* (Li et al., 2009; Lu et al., 2009; Nekrasov et al., 2009; von Numers et al., 2010; Farid et al., 2013). Therefore, it is unlikely that bioinformatics tools that predict the presence of N-glycosylated proteins in roots will be sufficient to identify the client N-glycosylated proteins that contribute to the Fe-dependent reduction in primary root growth under LPi conditions. Instead, a more promising approach would be a proteomic analysis aimed at experimentally detecting proteins adversely affected by the absence of calnexin.

## Material and Methods

### Plant lines and growth conditions

*Arabidopsis thaliana* seeds were surface sterilized and grown for 7 days on plates containing half-strength Murashige and Skoog (MS) medium without phosphate (Caisson Laboratories) supplemented with 75 μM or 1 mM KH_2_PO_4_ buffer (pH 5.8), 1% (w/v) sucrose, 0.7% (w/v) agarose, and 500 mg/L 2-(N-morpholino) ethanesulfonic acid (final pH 5.8). To induce different levels of phosphate and iron deficiency, ferrozine was added to the medium at a final concentration of 100 μM. Plants were grown vertically on plates at 22°C under a continuous light intensity of 100 μmol m^−2^ s^−1^.

Plants were also grown in soil or in a clay-based substrate (Seramis) irrigated with phosphate-free half-strength MS supplemented with KH_2_PO_4_ buffer, pH 5.8. The growth chamber conditions were 22°C and 60% relative humidity with a 16-h-light/8-h-dark photoperiod with 100 μE/m^2^ per s of white light.

All Arabidopsis lines used in this study are in the Col-0 background. A single *cnx1* (SALK_083600C) allele and two *cnx2* (SAIL_865_F08 and SAIL_580_H02) mutant alleles were identified from T-DNA insertional lines obtained from the European Arabidopsis Stock Center (NASC) (http://arabidopsis.info). Supplemental Table S1 lists the sources of all other lines used in this study. Plants overexpressing *PDR2* under the control of the CaMV35S promoter (Ticconi et al., 2009) as well as plants expressing the reporter construct cycB1::GUS (Colon-Carmona et al., 1999) were described previously.

### Phosphate quantification

Quantification of Pi was performed as previously described (Ames, 1966). Shoot or root material was placed in pure water, and at least three freeze-thaw cycles were applied to release the inorganic Pi, which was quantified via a molybdate assay using a standard curve.

### DNA constructs and gene expression analysis

PCR-generated fragments of the *CNX1* and *CNX2* genomic regions lacking stop codons and including the 1-kbp promoter regions were obtained using Phusion HF DNA polymerase (New England Biolabs), inserted into pENTR-2B, and recombined in pMDC107 to generate the GFP-tagged construct using Gateway technology. The binary vectors were introduced into Arabidopsis plants via *Agrobacterium tumefaciens*–mediated transformation using the floral dip method (Clough and Bent, 1998).

Total RNA was extracted from roots using an RNA Purification kit as described by the manufacturer (Promega), followed by DNase I treatment. cDNA was synthesized from 1 µg of RNA using M-MLV Reverse Transcriptase (Promega) and oligo d(T)_15_ following the manufacturer’s instructions. qPCR analysis was performed using SYBR Select Master Mix (Applied Biosystems) with primer pairs specific to genes of interest; *ACT2* was used for data normalization. The primer sequences are listed in Supplemental Table S2.

### Root measurements, microscopy, and staining procedures

Root length was measured using seedlings grown on vertically oriented plates. The plates were scanned on a flatbed scanner to produce image files suitable for quantitative analysis using ImageJ software (v1.44p).

Confocal microscopy was performed using a Zeiss LSM 880 confocal laser scanning microscope. Plant roots were treated with Clearsee solution and stained with calcofluor white (Ursache et al., 2018) to visualize cell walls. A line expressing the cycB1::GUS reporter was used to introgress the construct into the *cnx1-1 cnx2-2* double mutant background. Roots were stained for GUS activity as previously described (Lagarde et al., 1996). The tissues were vacuum infiltrated to enhance tissue penetration. Stained tissues were cleared in chloral hydrate solution (2.7 g/mL in 30% glycerol) and analyzed using a Leica DM5000B bright-field microscope.

Iron accumulation in seedlings was assayed by Perls-DAB staining as previously described (Müller et al., 2015). Briefly, seedlings were incubated in 4 mL of 2% (v/v) HCl and 2% (w/v) potassium ferrocyanide for 30 min. The samples were washed with water and incubated for 45 min in 4 mL of 10 mM NaN_3_ and 0.3% H_2_O_2_ in methanol. The samples were then washed with 100 mM Na-phosphate buffer (pH 7.4) and incubated for 30 min in the same buffer containing 0.025% (w/v) DAB and 0.005% (v/v) H_2_O_2_. Finally, the samples were washed twice with water, cleared with chloral hydrate (1 g/mL, 15% glycerol), and analyzed using an optical microscope.

### Immunoblot analysis

Proteins were extracted from homogenized plant material at 4°C in extraction buffer containing 10 mM phosphate buffer, pH 7.4, 300 mM sucrose, 150 mM NaCl, 5 mM EDTA, 5 mM EGTA, 1 mM DTT, 20 mM NaF, and 1× protease inhibitor (Roche EDTA Free Complete Mini Tablet) and sonicated for 10 min in an ice-cold water bath. Fifty micrograms of proteins were separated by SDS-PAGE and transferred to an Amersham Hybond-P PVDF membrane (GE Healthcare). The membrane was probed with rabbit polyclonal antibodies against maize calreticulin, which cross-reacts with both Arabidopsis calnexin and calreticulin (Persson et al., 2003), and goat anti-rabbit IgG-HRP (Santa Cruz Biotechnology) using Western Bright Sirius HRP substrate (Advansta). Signal intensity was measured using a GE Healthcare ImageQuant RT ECL Imager.

## Acknowledgments

The authors are grateful to Shuh-ichi Nishikawa (Niigata University, Japan) and Cyril Zipfel (University of Zurich, Switzerland) for seeds of the *bip* and *sdf2* mutants, respectively.

## Supplemental data

### Supplemental Figures

**Figure S1. Localization of *CNX1::CNX1-GFP* and *CNX2::CNX2-GFP* in the ER. (A)** Expression of *CNX1::CNX1-GFP* and *CNX2::CNX2-GFP* in roots tips of transgenic *cnx1-1 cnx2-2* plants. Bars = 10 µm. **(B)** Transient co-expression of *CaMV35S::CNX1-GFP* and *CaMV35S::CNX2-GFP* with the ER marker ER-RFP in tobacco leaves.

### Supplemental Tables

**Table S1**. List of mutants used in this study.

**Table S2**. Primer list.

