## Supplemental figure and tables for "Endoplasmic reticulum calnexins participate in the primary root growth response to phosphate deficiency"

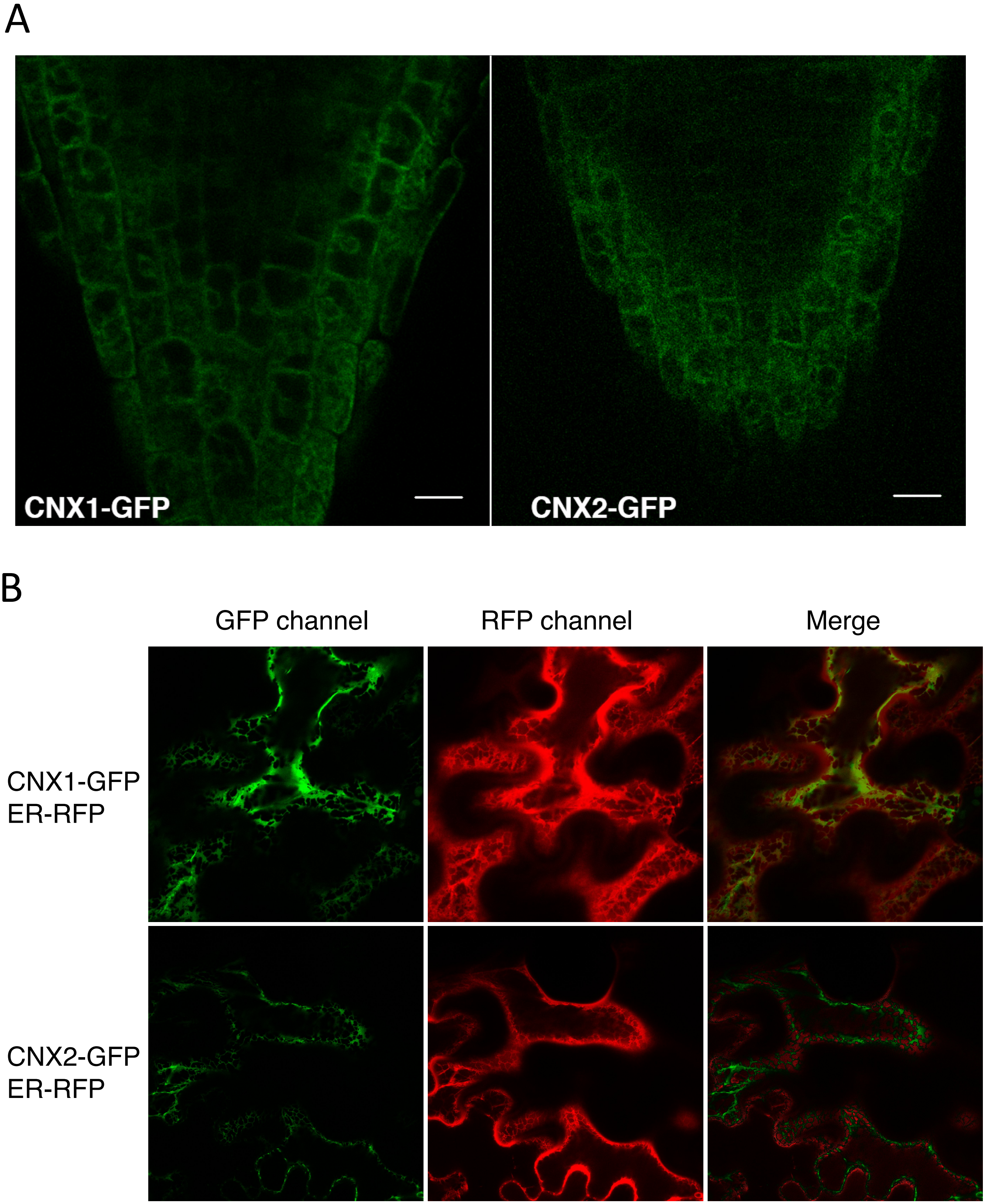


**Figure S1. Localization of *CNX1::CNX1-GFP* and *CNX2::CNX2-GFP* in the ER. (A)** Expression of *CNX1::CNX1-GFP* and *CNX2::CNX2-GFP* in roots tips of transgenic *cnx1-1 cnx2-2* plants. Bars = 10 µm. **(B)** Transient co-expression of *CaMV35S::CNX1-GFP* and *CaMV35S::CNX2-GFP* with the ER marker ER-RFP in tobacco leaves.

**Supplemental Table S1. List of mutants used in this work**

| Mutant | Gene code | Source | Function | Source |
|---|---|---|---|---|
| *cnx1-1* | AT5G61790 | SALK_083600 | Calnexin 1 | this work |
| *cnx2-2* | AT5G07340 | SAIL_865_F08 | Calnexin 2 | this work |
| *cnx2-3* | AT5G07340 | SAIL_580_H02 | Calnexin 2 | this work |
| *crt1-1 crt2-1* | AT1G56340, AT1G09210 | SALK_142821 SALK_099778 | Calreticulin 1, Calreticulin 2 | this work |
| *crt3-1* | AT1G08450 | SALK_051336 | Calreticulin 3 | this work |
| *alg3-1* | AT2G47760 | SALK_064006 | ER α-1,3-mannosyltransferase | Veit et al., Front Plant Sci 9, 1807 (2018) |
| *alg9a* | AT1G16900 | GABI_831D07 | ER α-1,2-mannosyltransferase | Veit et al., Front Plant Sci 9, 1807 (2018) |
| *alg10-1* | At5gG02410 | SAIL_515_F10 | ER α-1,2-glucosyltransferase | Farid et al., Plant J 68, 314 (2011) |
| *stt3a-2* | AT5G19690 | SALK_058814 | Oligosaccharyl transferase | Koiwa et al., Plant Cell 15, 2273 (2003) |
| *mns4 mns5* | AT5G43710, AT1G25720 | SALK_119093, GT_5_84786 | ER-a-mannosidase-like | Hüttner et al., Plant Cell 26(4), 1712 (2014). |
| *ebs1-6/uggt1-1* | AT1G71220 | WiscDsLox413-416K21 | UDP-glucose:glycoprotein glucosyltransferase | Jin et al., Mol Cell 26, 821 (2007) |
| *psl4* | At5g56360 | SALK_074006 | Beta-subunit of glucosidase II | this work |
| *bib1-4 bip3-1* | AT5G28540, AT1G09080 | CS856879 SALK_024133 | ER chaperone | Maruyama et al., Plant Cell Physiol 55, 801 (2014) |
| *bip2-1 bip3-1* | AT5G42020, AT1G09080 | CS842467 SALK_024133 | ER chaperone | Maruyama et al., Plant Cell Physiol 55, 801 (2014) |
| *sdf2-1* | AT2G25110 | T-DNA activation-tagging | complex with ERdj3B-BiP ER chaperones | Nekrasov et al., EMBO J 28, 3428 (2009) |
| *pdr2* | AT5G23630 | EMS mutant | P-type ATPase | Ticconi et al., Plant J 37, 801 (2004) |
| *lpr1-1* | AT1G23010 | SALK_016297 | Ferroxidase | Svistoonoff et al., Nature Genet 39, 792 (2007) |
| *lpr2-1* | AT1G71040 | SALK_091930 | Ferroxidase | Svistoonoff et al., Nature Genet 39, 792 (2007) |
| *lpr1-1 lpr2-1* | AT1G23010, AT1G71040 | SALK_016297, SALK_091930 | Ferroxidase | Svistoonoff et al., Nature Genet 39, 792 (2007) |

Supplementary Table S2. Primer list

| Gene ID | Gene name | Primer name | Sequence 5' - 3' |
| --- | --- | --- | --- |
| At5g61790 | CNX1 | qCNX1 Fw | CGCTGGATCGTTTCGAAGAA |
| | | qCNX1 Rv | CACACTCAAGTCCTTCCTGGAA |
| At5g07340 | CNX2 | qCNX2 Fw2 | AGTTCCCTCCTTCTGTTCCG |
| | | qCNX2 Rv2 | AGGAATCAACGGAGGCTCAA |
| At2G38940 | PHT1,4 | qPHT1,4 Fw | TTGCTCCTAATTTTCCTGATGCT |
| | | qPHT1,4 Rv | TGTGCCGGCCGAAATCT |
| At2G11810 | MGD3 | MGD3 qPCR Fw | AGAGGCCGGTTTAATGGAG |
| | | MGD3 qPCR Rv | CATCAGAGGATGCACGCTA |
| At1g42990 | bZIP60 | qbZIP60 Fw | CCGGCGGAGGATTTTCTTCA |
| | | qbZIP60 Rv | GCCAAATCAACGGAGCCAGA |
